# Are flexible school start times associated with higher academic grades? A 4-year longitudinal study

**DOI:** 10.1101/2021.07.29.452310

**Authors:** Anna M. Biller, Carmen Molenda, Fabian Obster, Giulia Zerbini, Christian Förtsch, Till Roenneberg, Eva C. Winnebeck

## Abstract

The mismatch between teenagers’ late sleep phase and early school start time results in acute and chronic sleep reductions. This is not only harmful for students’ learning in the short-term but may impact on students’ career prospects and widen social inequalities. Delaying school start times has been shown to improve sleep but whether this translates to better achievement is unresolved. The current evidence is limited due to a plethora of outcome measures and the many factors influencing sleep and grade/score trajectories. Here, we studied whether 0.5-1.5 years of exposure to a *flexible* school start system, with the *daily* choice of an 8AM or 8:50AM-start (intervention), allowed secondary school students (n=63-157, 14-19 years) to improve their quarterly school grades in a 4-year longitudinal pre-post design. We investigated whether sleep, changes in sleep or frequency of later starts predicted grade improvements in the flexible system. Our mixed model regressions with 5,111-16,724 official grades as outcomes did not indicate meaningful grade improvements in the flexible system *per se* or with previously observed sleep variables (nor their changes) – the covariates academic quarter, discipline and grade level had a greater, more systematic effect in our sample. Importantly, this finding does not preclude improvements in learning and cognition in our sample. However, at the ‘dose’ received here, intermittent sleep benefits did not obviously translate into detectable grade changes, which is in line with several other studies and highlights that grades are suboptimal to evaluate timetabling interventions despite their importance for future success.

**Significance statement:** Early school start times worldwide clash with teenagers’ delayed sleep-wake rhythms. This mismatch results in sleep restrictions below healthy amounts, which compromises health and performance and further aggravates social disparities. Since adequate sleep is important for learning and concentration, there is the strong expectation that counteracting sleep deprivation with delayed school starts results in better academic achievement. We add important high-resolution, longitudinal data to this unresolved scientific debate. When controlling for confounders, our results do not support that improved sleep leads to grade changes within 1.5 years in a flexible start system. While grades are suboptimal to measure later school start effects on performance, they nevertheless open doors to higher education worldwide and thus determine future trajectories.

## Introduction

During adolescence, teenagers undergo a plethora of biological and socially-driven developments that also influence their sleep-wake behaviour [1–4]. Their internal phase (chronotype) delays progressively with age until around 21 [5], while sleep pressure (the homeostatic load) likely accumulates more slowly across the day compared to adults [6,7]. This predisposes teenagers, more than younger children or adults, to delay into late evening hours thereby also delaying their sleep timing. Early school start times cut teenagers’ sleep artificially short in the morning, forcing them to get up before they reach healthy amounts of 8-10 hours of night-time sleep on schooldays. On weekends, teenagers sleep not only longer but also later which better suits their delayed circadian clock. This sleep timing difference between school and free days is called “social jetlag”, since it is a constant jetlag situation induced by social schedules [8,9]. These widespread sleep restrictions are not only connected to compromised health and well-being [10–14], but also reduced cognition (e.g. constructive [15] and creative thinking skills [16,17] or verbal fluency [18,19]) and decreased academic performance [20].

There is now substantial amount of evidence that delayed school start times help to increase sleep durations towards more healthy amounts, at least in the short-term [e.g. 21–24]. However, it is less clear whether the increased sleep durations and better sleep quality also translate to better learning in the same students. This could reasonably be expected, given the prominent role of sleep for memory, sustained attention and concentration [e.g. 25–27].

Thus, several studies previously tried to evaluate the effects of delayed school start times on grades or scores. These studies yielded very mixed results, probably due to differences in study designs, interventions, exposure times and outcome measures [28]. However, all interventions assessed were static changes in school start times whereas the possibility to make school start times flexible has been largely overlooked.

Here, we studied whether a *flexible* school start system, as implemented in a secondary school in Germany [29,30], and concurrent changes in sleep [29,30], were associated with changes in grades in multiple academic disciplines. The flexible system entailed that the school changed from a permanent fixed start at mainly 8AM to a *flexible* school start that allowed senior students to choose *daily* whether to attend school at 8AM or skip the first class and start at 8:50AM. To examine effects of this new system on academic grades in detail, we analysed students’ quarterly grades from 12 academic school subjects across 4 years. With 2.5 years of data prior and up to 1.5 years after the introduction of the flexible school start, we could control for some important confounders and address trends and complex interactions which started long before the system was changed. In addition, we also used longitudinal sleep data which we had previously collected in these students [29,30] to predict students’ quarterly grades by means of linear mixed regression models.

## Methods and Materials

### Study Site and the Flexible School Start System

The study took place at the Gymnasium Alsdorf (50° 53’ N, 6° 10’ E), a secondary school in the West of Germany. This particular school offers daily self-study periods during which students work through a personal 5-week curriculum with a teacher and on a subject of their own choice (so-called “Dalton system”).

On February 1^st^, 2016, the school changed permanently from a fixed start (“conventional system”) to a flexible start (“flexible system”) for 10-12^th^ graders (senior students). In the conventional system, school started at 8AM on most days. On a median of 1 day/week (depending on students’ individual timetables), however, school started with the second period at 8:50AM. In the flexible system, the first period (lasting 08:00-08:45AM) was made optional for senior students to attend. Senior students could thus choose daily whether to start at 8AM with the first self-study period or skip it and start at 08:50AM instead (called “9AM” here for convenience). On a median of 1 day/fortnight, students also had a scheduled free second period (08:50-09:50AM), i.e. the chance to start at 10:15AM (“>9AM”). Given the low occurrence rate of >9AM-starts, we did not distinguish between frequencies of 9AM-starts and >9AM-starts in our analyses. For more information on the flexible system, please refer to [29].

### Study Design

Official academic grades were obtained from the school registry for students that took part in our first wave in 2016 [29] and the second wave in 2017 [30]. While grades were provided retrospectively at the end of the schoolyear of wave 2 for the past 4 years, sleep data were collected longitudinally in both waves (Fig. 1): Wave 1 consisted of baseline sleep diary data collection (=t0) covering 3 weeks in January, 2016 (Jan 8^th^ to 31^st^) in the conventional system, followed by sleep diary data collection for 6 weeks (Feb 1^st^ to Mar 14^th^) in the flexible system right after its introduction on Feb 1^st^, 2016 (=t1). Wave 2 covered the matching photoperiod and time of t1, lasting again 6 weeks (Feb 2^nd^ to Mar 20^th^, 2017, = t2). No second baseline just before t2 was carried out since the school had remained in the flexible system. We excluded any sleep diary entries during the carnival holiday periods between February 4^th^-9^th^, 2016 and February 23^rd^-28^th^, 2017 from the analyses.

**Figure 1.**
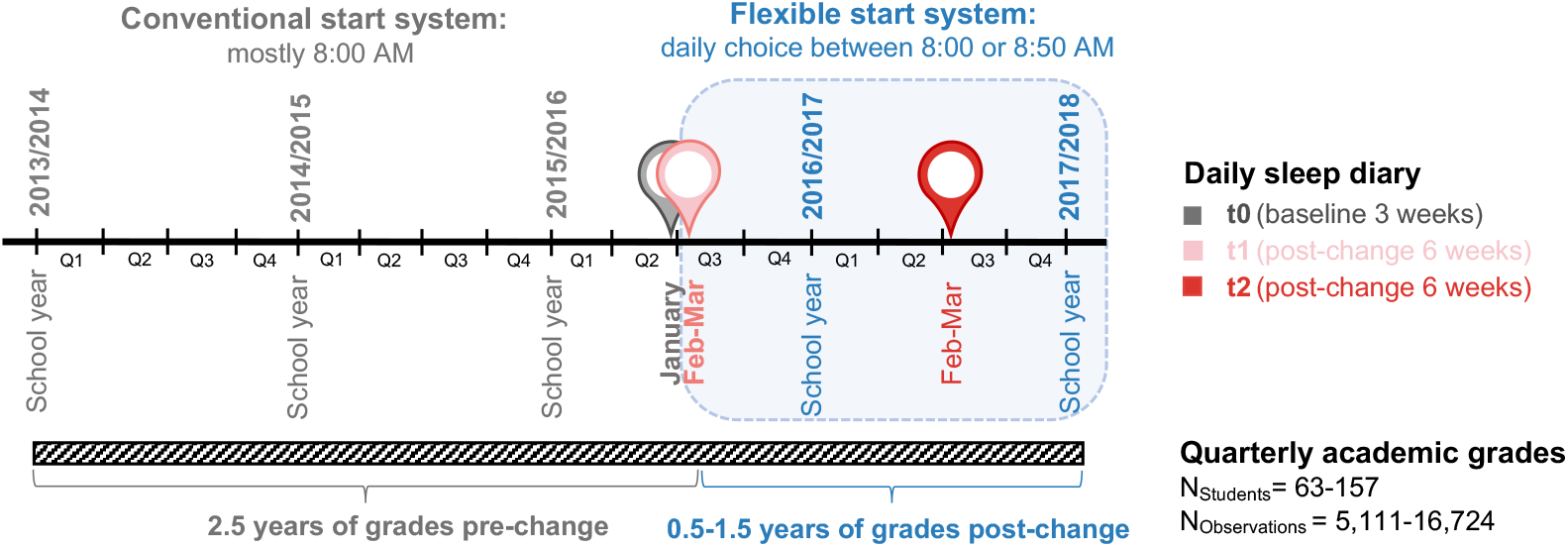
Study design and outcome measures. Schematic of longitudinal study design including quarterly academic grades from up to 2.5 years prior to and up to 1.5 years after the introduction of the flexible system. The same students had also provided daily sleep diary data in two waves (one baseline assessment in the conventional system and 2 time points in the flexible system as described previously [30].

### Participants

Written informed consent was obtained from all participants (or their parents/guardians if <18y). The study was conducted according to the Declaration of Helsinki and approved by the school board, the parent-teacher association, the school’s student association and the ethics committee of the Medical Faculty of the LMU Munich (#774-16). We used opportunity sampling without specific exclusion criteria. For response and attrition rates and filter criteria of sleep diary data and cohorts please refer to [30]. Academic grades were provided by the school registry from students that took part at any time during our study (t0 through t2). All included students were granted promotion to the next grade level during the study period.

## Outcome measures

### Sleep Diary

We used a daily sleep diary (provided online via LimeSurvey.org) based on the μMCTQ [31]. Students provided sleep onset (note: not bedtime) and offset (wake time) of their past night’s sleep, the type of day they woke up (schoolday or free day), and when they started school (8AM, 9AM or >9AM). The questionnaire did not cover any naps during the day. For detailed sleep diary descriptions, please see [29,30].

### Academic Grades

From the school registry, we obtained official quarterly grades awarded to participating students between the school year 2013/2014 through to 2016/2017. Of the 170 students from both waves qualifying for analysis (i.e. the cohorts previously described and used for sleep analyses; see [30]), 13 students had grades missing, thus resulting in a maximum sample of 157 students for the grade analyses. For the majority of these students (62%), grade data span 2.5 years in the conventional and 1.5 years in the flexible system; for those in grade level 10 at wave 2 (18%), it was 3 and 1 years, and for those at grade level 12 at wave 1 (15%) it was 2.5 and 0.5 years. The grades were provided for all academic subjects taken by a student, of which we included only 12 subjects in our analyses that most students took and assigned them to one of three disciplines: Sciences (Biology, Chemistry, Maths, Physics, Natural Sciences), Social Sciences (Geography, History), and Languages (English, German, Spanish, French, Latin). Provided grades were averages per academic quarter per academic subject over a mixture of written and oral examinations, course work and participation in class.

The school year lasted from the end of August to mid-July divided into the following quarters: quarter 1 until end of October, quarter 2 until third week of January, quarter 3 until third week of April and quarter 4 until first week of July.

In grade levels 7-10, the grading scale ranged from 1 (best) to 6 (worst) with grades ≤4 considered passing grades. This scale was additionally broken down into plus (+) and minus (-) for all but grade 6. In grade levels 11 and 12, the scale ranged from 0 (worst) to 15 (best) with ≥4 considered passing. Both scales were combined by transforming the 1-6 scale to a 0-15 scale based on its finer plus/minus system and then transformed to a more universal 0%-100% scale.

## Data Analysis

Analyses were performed in SPSS Statistics (IBM, versions 24 and 25) and R (versions 3.6.1 and 3.6.3) using R studio (versions 1.1.463, 1.2.1335 and 1.2.5042). Graphs were produced using the r-package *ggplot2* [32].

### Sleep Data

Daily sleep data from dairies were aggregated and taken from t2 if available (else from t1). From these aggregates, we derived the following variables as per equations below: average daily sleep duration during the week (SD_week_), chronotype as midsleep on free days (MSF) corrected for oversleep (MSF_sc_), social jetlag (SJL) as midsleep on free days (MSF) minus midsleep on work/school days (MSW), and frequency of ≥9AM-starts. For the linear mixed models 3a-d, we additionally calculated the absolute differences between t0 and t1 (i.e., from baseline to the flexible system during wave 1) for SD_week_, MSF_sc_ and SJL (X change).

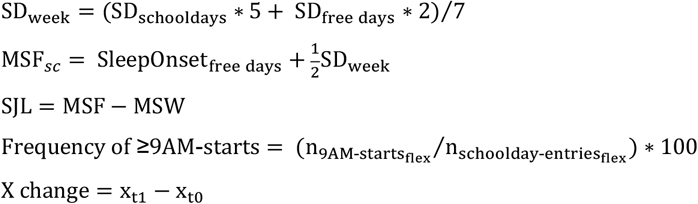

### Statistical analyses

Unless indicated otherwise, descriptive statistics are reported as mean ± standard deviation and test statistics are abbreviated as follows: *t*, t-test; Pearson correlation; *rho*, Spearman rank correlation; b, unstandardized coefficient of linear regression or linear mixed models; b_flex*change_, unstandardized coefficient of the interaction of linear mixed models; *p*, significance level. The alpha-level was set to *p*<0.05 for all statistical analyses. All data were tested on normality (histograms, QQ plots, Shapiro-Wilk’s test) and sphericity.

For simple grade analyses comparing accumulative grade point averages in the conventional versus the flexible system, a two-sided paired t-test was used. To this end, the grade point average was calculated as the mean grade across all subjects before or after the start-system change for each student. For more sophisticated grade analyses, we used linear mixed-effects regression models (*lme4* and *lmer test* package [33,34] in R). In total, 4 different models (plus model variations) were calculated to answer different questions based on different fixed effects, interaction terms and subcohorts (see overview Tab. 1). Student ID was added as random effect to all models to incorporate unsystematic differences between individuals. In all models, the outcome (dependent variable) was quarterly grades per discipline per student; the fixed effects (independent variables) were system (conventional/flexible), gender (female/male), grade level (7-12), academic quarter (1-4), and academic discipline (Sciences/Social Sciences/Languages), all entered as categorical variables. Model 1 additionally included interaction terms between discipline and gender to assess general grade influences, model 2 included interaction terms between school start system and gender, and system and discipline to assess system effects per discipline and gender. In models 3, we included one of the aggregated sleep-change variables (see equation above; mean-centred) as additional fixed effects, each in interaction with system (conventional/flexible): chronotype change (model 3a), sleep duration on schooldays change (model 3b), social jetlag change (model 3c) or frequency of ≥9AM-starts (model 3d). In model 4, we instead included the absolute value of chronotype, sleep duration on schooldays, social jetlag, and frequency of ≥9AM-starts for the flexible system only (from t2 if available, else from t1 to maximize sample size). Since chronotype, sleep duration on schooldays, social jetlag, and frequency of ≥9AM-starts were prone to collinearity, we first assessed their correlations before adding them into the models (Fig. S1). Only chronotype and social jetlag were highly correlated (rho = 0.65, p<0.001; Fig. S1), and results from models including just one of these variables each (4a-d) were essentially similar to model 4e which included all sleep variables together (Tab. S4). The variance inflation factor (*car* package in R [35]) also indicated no problematic collinearity for model 4e. Marginal means of model estimates were calculated using *emmeans* in R [36] for models where interactions were significant. All linear mixed models were visualised in tables using the *sjPlot* and *sjmisc* packages [37,38] and in figures as marginal means via the *ggeffects* package [39] in R. Simple contrast results from interactions in linear mixed models were averaged over the levels of system or gender (depending on the model), grade level, and quarter; degrees of freedom method used was Kenward-Rogers. Pairwise comparisons were adjusted with Tukey method.

**Table 1.** Overview of linear mixed model analyses on official, quarterly grades. Four different models (and several model variations) were calculated, each with a different aim and including appropriate predictors (fixed effects) and interaction terms. All models included ID as a random intercept to incorporate random inter-individual differences. Abbreviations: conv, conventional school start system; flex, flexible school start system.

|  | Model 1 | Model 2 | Model 3a-d | Model 4a-e |
| --- | --- | --- | --- | --- |
| **Outcome** | Official grades (per quarter and academic subject) | Official grades (per quarter and academic subject) | Official grades (per quarter and academic subject) | Official grades (per quarter and academic subject) |
| **Aim** | General effects | System effects | Effects of sleep changes | Effects of sleep in flexible system only |
| **Fixed effects** | System (conv/flex)<br>Gender<br>Grade level<br>Academic quarter<br>Academic discipline | System (conv/flex)<br>Gender<br>Grade level<br>Academic quarter<br>Academic discipline | System (conv/flex)<br>Gender<br>Grade level<br>Academic quarter<br>Academic discipline<br>Change<sup>a</sup> in...<br>a. ... chronotype<br>b. ... sleep duration<br>c. ... social jetlag<br>d. 9AM-use<sup>b</sup> | -<br>Gender<br>Grade level<br>Academic quarter<br>Academic discipline<br>-<br>a. Chronotype<sup>c</sup><br>b. Sleep duration<sup>c,d</sup><br>c. Social jetlag<sup>c</sup><br>d. 9AM-use<br>e. all of the above |
| **Interactions** | Gender *<br>Academic discipline | System *<br>Academic discipline<br>System *<br>Gender | System *<br>Chronotype change /<br>Sleep duration change /<br>Social jetlag change /<br>≥9AM-use | - |
| **Random intercept** | ID | ID | ID | ID |
| **Sample** | Waves 1 & 2 | Waves 1 & 2 | Wave 1 only | Waves 1 & 2 |
| **N** | 157 | 157 | 63 | 129 |
| **Number of observations** | 16724 | 16724 | 6683 | 5111 |
| **Data span** | 4 years:<br>2.5 y conv & 1.5 y flex | 4 years:<br>2.5 y conv & 1.5 y flex | 4 years:<br>2.5 y conv & 1.5 y flex | 1.5 years:<br>only flex |

<sup>a</sup> Change refers to the absolute difference between the respective variable at t1 minus t0 (baseline). Positive values indicate higher numbers at t1.
<sup>b</sup> Since the exact frequency of 9AM-starts during baseline (t0) is not known, 9AM-use was added as an absolute value rather than the change from t0 to t1. Students attended school at ≥9AM at a median of 1 day per week in the conventional system.
<sup>c</sup> From t2 if possible, else from t1.
<sup>d</sup> Duration on schoolday.

## Results

In this study, we investigated whether a *flexible* school start system and concurrent changes in sleep (as previously described here [29,30]) were associated with changes in academic grades. During a first wave of sleep assessment [29], the studied secondary school had changed from a conventional start system (mostly starting at 8:00AM; baseline=t0) to a flexible start system with a daily choice between 8:00 or 8:50AM (=t1; Fig. 1). The school has since maintained this system allowing for a second wave of sleep assessment after exactly one year (t2; [30]).

For the current study, we included quarterly grades of 63-157 students from these two waves irrespective of time of participation (i.e. t0/t1, t2 or all time points) (Fig. 1). The sample size varies depending on the analysis question and thus with the respective regression model calculated (see Tab. 1). The majority of included students were females (63%-68%), were in grade levels 10 or 11 (but levels 9 and 12 were also included), and used the late-start option (“≥9AM-use”) on about 24%-28% of all recorded schooldays (see Tab. 2 for more cohort characteristics). In total, we analysed 5,111-16,724 grades (on average 107 individual grades per person) that students received in 12 academic subjects over 2.5 years in the conventional system and 0.5 to 1.5 years in the flexible system (Fig. 1). Grades were provided by the school registry and transformed to a 0%-100% scale and labelled (not aggregated) as Languages, Sciences, or Social Sciences. Median grades were 53%-60% in Languages, 60% in Sciences, and 60%-67% in Social Sciences (Tab. 2). To complement the analyses, we also used several of the existing sleep variables (chronotype expressed as MSF_sc_, sleep duration, social jetlag) in the flexible system and their respective change (delta) from baseline (t0) to the flexible system (t1) as well as the frequency of ≥9AM-use in several model specifications (Tab. 1 and Tab. 2).

**Table 2.** Composition of study cohorts. Displayed are cohort characteristics after standard filter criteria. Grades were provided by the school registry. Grade levels are taken from t1. Abbreviations: n, number of individuals; SD, standard deviation; IQR, interquartile range; conv., conventional.

|  |  | Sample models 1 and 2 | Sample model 3 | Sample model 4 |
| --- | --- | --- | --- | --- |
| **Participants** |  |  |  |  |
| Total | n | 157 | 63 | 129 |
| Females | n (%) | 109 (69%) | 40 (63%) | 88 (68%) |
| Grade level | n (%) per level<br>9<sup>th</sup>/ 10<sup>th</sup>/11<sup>th</sup>/12<sup>th</sup> | 29/50/52/26<br>(18/32/33/17%) | 0/25/21/17<br>(0/40/33/27%) | 24/43/45/17<br>(19/33/35/13%) |
| **Academic grades per discipline on a scale from 0% (worst) – 100% (best)** |  |  |  |  |
| Languages | median<br>(IQR) | 53%<br>(40-73) | 53%<br>(47-70) | 60%<br>(46-73) |
| Sciences | median<br>(IQR) | 60%<br>(47-73) | 60%<br>(47-73) | 60 %<br>(46-73) |
| Social Sciences | median<br>(IQR) | 60%<br>(53-73) | 60%<br>(53-73) | 67%<br>(53-73) |
| **Sleep variables** |  |  |  |  |
| | | | Change ($\Delta$ t1-t0) | t2 if available,<br>else t1 |
| Chronotype<br>(MSF<sub>sc</sub>; time in h) | mean<br>(SD, range) | not used in model | 0.2<br>(0.7, -1.6-2.2) | 4.7<br>(1.0, 2.2-8.6) |
| Social jetlag (h) | mean<br>(SD, range) | not used in model | -0.3<br>(0.7, -2.0-1.9) | 2.0<br>(0.8, 0.5-6.0) |
| Sleep duration (h) | mean<br>(SD, range) | not used in model | 0.1<br>(0.5, -1.3-1.1) | 7.2<br>(0.8, 5.2-9.0) |
| **Frequency of attending school later** |  |  |  |  |
| ≥9AM-use | median<br>(IQR) | not used in model | 24%<br>(14-46) | 28%<br>(10-52) |

### School start system showed no systematic effect on academic grades overall

First, we investigated whether the flexible system allowed students to increase their grades without considering sleep variables. At first sight, a simple comparison of overall grades yielded a small but statistically significant improvement in grade point average from 58.2% (±2.1 SD) in the conventional to 59.6% (±2.0 SD) in the flexible system (Fig. 2a; t[154]=-2.15, p=0.033, d_z_=0.173). However, attributing this improvement to the flexible system is likely unwarranted. As outlined in the introduction, grades are influenced by a multitude of factors, thus comparisons that do not account for these can be misleading. We therefore applied linear mixed-effects regression models to adjust for potential confounders and including random intercept for ID to account for inter-individual differences (Tab. 1). When incorporating gender, grade level (i.e. indirectly age), academic quarter, and discipline in addition to school start system in the analysis, the flexible system showed no systematic relationship with students’ grades (Fig. 2b; b= -0.10, p=0.815, Model 1, Tab. S1), hence the flexible system was not associated with students receiving better or worse grades overall in our sample (n_students_=157).

**Figure 2.**
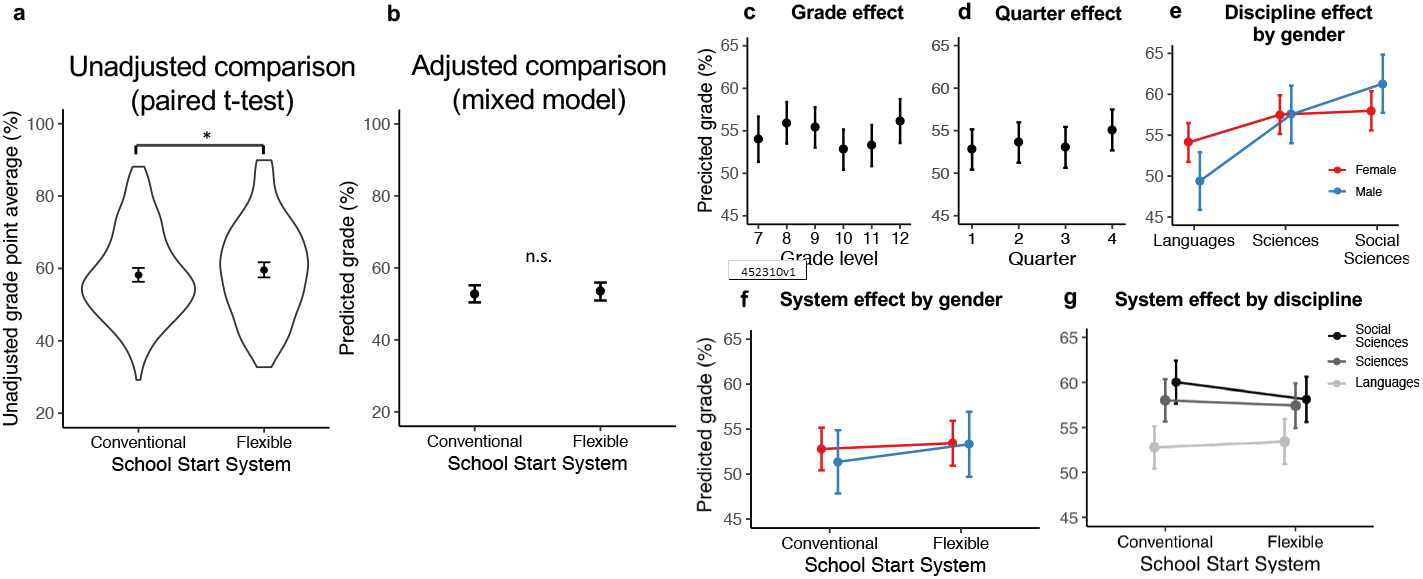
Longitudinal analysis of official quarterly grades - effects of school start system and general predictors. Quarterly grades (0%-100%) from 12 academic subjects of 3 disciplines for 4 years i.e., for most students this was 2.5 years before and 1.5 years after the flexible school start was introduced (n=157 students; 16,724 grades; 107 grades per student on average). **a**, Simple, unadjusted comparison of average grades across all disciplines in the conventional and the flexible school start system via paired t-test (n_ID_=157). Shown are mean and 95% CI within the raw data distribution (violin plots). The apparent grade improvement in the flexible system was not confirmed in linear mixed models. **b-g**, Visualization of mixed-model-determined influences on grades. Plots show marginal means from models 1 and 2 (Tab. S1), i.e. the estimated grade and 95% CI for the reference category (female student, class level 10, quarter 1, languages, conventional system). Statistical significance is indicated in (b), results of more complex cases can be found in the text and Tab. S1 and Tab. S2. **b**, Effect of school start system (model 1). **c**, Effect of grade level (model 1). **d**, Effect of academic quarter (model 1). **e**, Effect of academic discipline by gender (model 1). **f**, Effect of school start system by gender (model 2). **g**, Effect of school start system by academic discipline (model 2).

### Grades varied systematically with grade level, academic quarter, discipline and gender

But what drives better grades in the unadjusted comparison if not the flexible system itself? The same factors that we adjusted for in the regressions also stood out as major predictors (Model 1, Tab. S1, n_students_=157): Students in 12^th^ grade (the last school year) did consistently better compared to their peers across all other grade levels – a sort of “leavers effect” that has already been observed before (Fig. 2c, b=3.44, p<0.001) [40]. Moreover, we found that students enjoyed a bump in grades in the last quarter of the school year with an estimated improvement of 2.3 percentage points compared to the first quarter (Fig. 2d; b=2.34, p<0.001). The combination of these two effects might explain the statistically significant improvement observed in the unadjusted comparison: the flexible system replaced the conventional system mid-year between quarter 2 and 3, so quarter 4 and higher grade levels were overrepresented in the flexible system, which the t-test could not account for.

The mixed models also revealed other strong systematic influences on grades in our sample. Firstly, we observed a clear difference between the disciplines: students performed generally best in Social Sciences, followed by Sciences and then Languages (Model 1, Tab. S1). Post-hoc tests (Fig. 2e) showed that these differences were highly significant for both genders (all p<0.001; post-hoc to Model 1, Tab. S2), except for girls’ grades in Sciences and Social Sciences, which were indistinguishable (b=-0.47, p=0.3895; post-hoc to Model 1, Tab. S2).

Female gender has been reported as another driving force for higher grades [41]. However, girls in our sample did not outperform boys overall (Model 2, Tab. S1 and Tab. S2). Girls were significantly better in Languages (Fig. 2e; b=4.72, p=0.0284; post-hoc to Model 1, Tab. S2), while boys surpassed them in the Social Sciences (b=-3.31, p=0.1269; post-hoc to Model 1, Tab. S2), and both genders did equally well in Sciences (b=-0.00, p=0.9915; post-hoc to Model 1, Tab. S2).

### The flexible system was linked with subtle improvements in Languages and subtle drops in Social Sciences grades

Although we did not find evidence that the flexible system was linked with better grades overall (Model 1, see above), the flexible system might be linked with grade improvements in certain disciplines and genders. To assess this, we looked at the interaction between i) school start system and discipline, as well as ii) school start system and gender in a second model (Model 2; Tab. S1, n_students_=157). Neither females nor males significantly improved their overall grades from the conventional to the flexible system (Fig. 2f; post-hoc to Model 2, Tab. S2). In terms of discipline effects, we found that grades in Social Sciences slightly dropped (b=1.26, p=0.0384; post-hoc to Model 2, Tab. S2), Science grades remained unchanged (b=-0.07, p=0.8849; post-hoc to Model 2, Tab. S2), and Language grades slightly improved (b=-1.30, p=0.0168, post-hoc to Model 2, Tab. S2) in the flexible system. Notably, these changes were subtle but reduced the grade differences between the academic disciplines (Fig. 2g, Tab. S2). These small changes in opposite directions likely explain the absence of a net effect of the flexible system on overall grades.

### Improvements in chronotype, sleep duration, and social jetlag did not systematically improve grades

What was the role of sleep parameters on grade developments? We speculated that students who showed greater improvements in the flexible system (i.e., advanced chronotype, lengthened sleep duration, and lowered social jetlag) also received better grades in the flexible system. Thus, we computed changes in sleep from t0 (baseline) to t1 based on the subpopulation of students with sleep parameters during these time points (n=63). Adding these parameters separately into a third model (Models 3a-c, Tab. S3, Fig. 3a), we found that neither changes in chronotype (flex*chronotype change: b=0.10, p=0.845) nor sleep duration (flex*sleep duration change: b= -0.77, p=0.352) were systematically associated with changes in grades. Surprisingly, however, students who increased their social jetlag in the flexible system obtained slightly better grades in the flexible system (flex*social jetlag change: b=1.28, p=0.027), which was contrary to our hypothesis. Therefore, our analyses in this subsample suggest that sleep improvements experienced immediately after transitioning to the flexible system did not result in detectable higher academic achievement.

**Figure 3.**
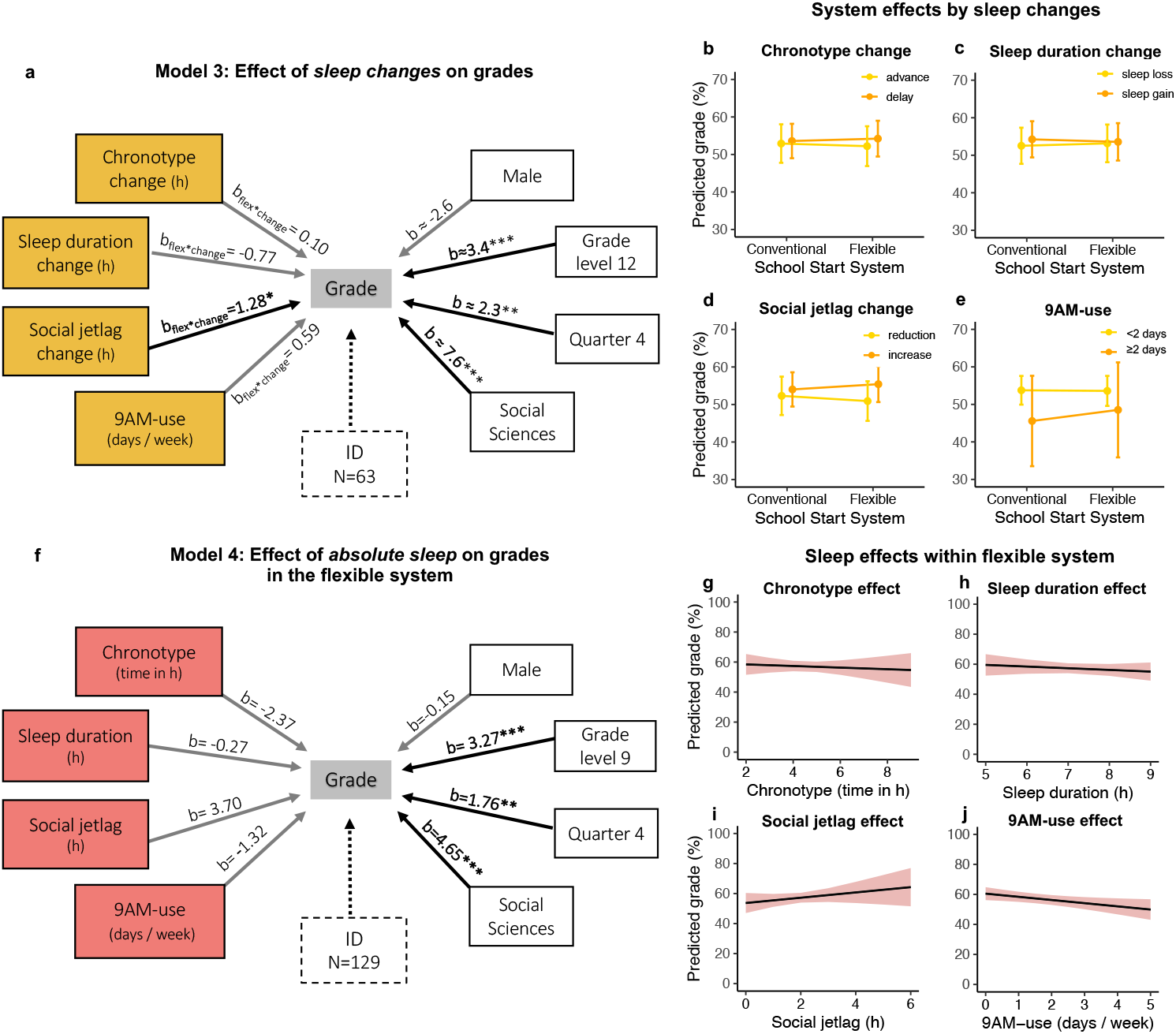
Longitudinal analysis of official quarterly grades - effects of sleep and 9AM-use. Results from linear mixed model analyses of quarterly grades (0-100%) considering sleep variables and the frequency of ≥9AM-starts (see Tab. 2 for sample descriptions). **a**,**f**, Schematic of the structure and results from models 3 and 4 (Tab. S3 and Tab. S4) showing the outcome, official quarterly grades (center), all predictors (black-framed boxes), the statistical significance of their effect (arrows; black: p<0.05, grey: p≥0.05), the unstandardized regression coefficients (b-values) and ID as random intercept (dashed box). General predictors (white) are categorical variables, so the levels with the highest impact are shown compared to their reference (female, grade level 10, quarter 1, languages). b-values are approximate in **a**, indicated by ≈, as representing results from models 3a-d. **a**, Effect of changes in sleep and of 9AM-use on grade improvements from the conventional to the flexible system. Summarized results from models 3a-d (n_ID_=63; Tab. S3) where each model variation included a different yellow predictor in interaction with school start system (conventional/flexible; b_flex*change_) to model effects of sleep changes on grade changes. **b-e**, Visualization of the yellow interaction effects from (a) via marginal means, i.e. grade estimates and 95% CI for the reference (female student, class level 10, quarter 1, languages) and categorical splits in the continuous sleep change variables to facilitate display. The effect of school start system on grades by **b**, chronotype change (advance/delay), **c**, sleep duration change (sleep loss/sleep gain), **d**, social jetlag change (reduction/increase) from the conventional to the flexible system, and by **e**, the frequency of 9AM-use (<2days/≥2 days) in the flexible system. **f**, Effect of absolute sleep characteristics on grades in the flexible system. Summarized results from model 4e (n_ID_=129; Tab. S4) predicting grades only for the flexible system, i.e., 1.5 years post-change, including the red sleep predictors in one common model after running separate models (4a-d) to check for collinearity. **g-j**, Visualization of the red effects from (f) via marginal means, i.e. grade estimates and 95% CI for the reference (female student, class level 10, quarter 1, languages). *, p<0.05; **, p<0.01; ***, p<0.001.

If not linked to sleep improvements, were grades nonetheless linked with the choice of more later school starts? The results of Model 3d calculated to answer this question suggests that higher 9AM-use was associated with worse grades in the conventional system (b=-3.04, p=0.015), a link reversed partly – albeit not significantly – in the flexible system (flex*9AM-use: b=0.59, p=0.101; Fig. 3a and Tab. S3). Hence, students who made high use of the late-start option in the flexible system were predominantly the lower-achievers, but they tended to benefit at least slightly from more later starts.

### No systematic effects of chronotype, social jetlag and sleep duration on grades

Lastly, we investigated if we could find absolute effects of sleep variables (chronotype, social jetlag, sleep duration) and ≥9AM-use on grades in the flexible system (Model 4, n=129 students). In contrast to what we had expected, none of the sleep parameters showed any significant link with grades, independent of whether they were added separately in the model (Model 4a-d) or together (Model 4e; Fig. 3f, Tab. S4). Our results thus indicate that late chronotypes in our sample were not worse off compared to their early peers and that longer sleep duration in the flexible system did not improve grades received in the flexible system. Similarly, social jetlag did not hamper grades to such a degree that we could detect an effect. Furthermore, we found that per every additional day a student chose to go to school later, grade estimates decreased by 1.52 (p=0.275, Tab. S4 Model 4e; for the single model 4d: b=-2.14, p=0.047). Although the interpretation is slightly different, this result tallies with the above finding: At first sight, it looks as if attending school later more often would prevent students from getting better grades, but we argue that most likely it is the other way around; students who receive worse grades also liked to attend school later more often when they had the chance to do so. On the whole, we could not show that chronotype and social jetlag negatively influenced grades, and it seemed as if mainly lower achieving students in our sample liked to use the ≥9AM option.

## Discussion

Adolescence is a decisive time in life for teenagers around the world. Teenagers undergo many cognitive, emotional and brain structural changes that also shape their risk-taking behaviour, learning capacities and motivation to attend school [42,43]. A prominent change also occurs in their daily sleep-wake behaviour: teenagers tend to phase-delay their sleep-wake behaviour, which essentially means that they become night-owls [5,44–48]. This delayed phase, however, clashes with early school starts seen across many countries, thus cutting sleep short in the morning hours during the school week. Apart from many other negative (health) consequences [10,11,49–52], short and low-quality sleep as well as sleepiness likely influences academic success thus shaping future career trajectories [20]. Since sleep restrictions and poor sleep habits are more severe in minority groups and disadvantaged students [53,54], addressing this problem is key to closing the achievement gap between social groups. However, the evidence is not conclusive whether delayed school start times can ameliorate this pressing health and performance problem. Many previous studies suffer from study design limitations, outcome variables are not comparable, and long-term studies that track individuals over time are rare [28]. Here, we studied whether a novel timetabling system – a daily chosen flexible school start – has the potential to improve academic grades via improved sleep.

In our study, we found that the flexible system was only associated with higher grades at first sight. When not adjusting for confounding factors, we observed a small improvement of grades in the flexible system, which would be in line with some previous studies [e.g., 55,56]. However, we argue that such simple pre-post analysis of aggregated grades is not suited to answer this complex question – although this has frequently been done using cross-sectional data. Studies on grades that performed proficient analyses, such as mixed regression models [57], quantile regression models [58] or difference-in-difference approaches [59–61] accounting for available confounders provided mixed results and mostly small effect sizes (for a systematic review see [28]). Nonetheless, positive effects of delayed school start times on academic achievement have been widely proclaimed – bound to raise falsely high expectations in parents and teachers. When we considered grade level, discipline and quarter in mixed model analyses, we found that the flexible system was clearly not associated with overall grade improvements except for subtle increases in Languages and subtle decreases in Social Sciences. In fact, the “confounders” weighed much stronger in our sample than any school start system effect on individual disciplines: graduating students did constantly better, highest grades were given in the final quarter of the year, and students were most successful in Social Sciences. Furthermore, the interplay between gender, discipline and school start system on grades is complex. Importantly, we also did not find any expected relationships between chronotype, social jetlag, or sleep duration with grades in our sample. Neither changes in these sleep parameters from the conventional to the flexible system nor their absolute values in the flexible system showed any link with grades - except for changes in social jetlag. Surprisingly, an increase in social jetlag, not a decrease, in the flexible system was predictive of higher grades in the flexible system. We have not been able to identify obvious explanations for this finding in exploratory analyses, except for the fact that weekend sleep was much more variant and backed by fewer data points than schoolday sleep, pointing towards a potential chance finding. A likely explanation for our null-finding for the other sleep parameters is a possible lack of power in our sample of 157 students (even though we have >16,000 longitudinal grades) given the small effect sizes previously identified (ranging around <0.1 SD; see [28]). A second possibility is that the time frame students were exposed to the new system was too short (exposure length) or that the delay was too little or infrequent (dose) in our study. Furthermore, sleep variables obtained at discrete study points might not be reflective of sleep during the other academic quarters or years. Thus, we cannot preclude that we missed a subtle effect in our sample but any such effect is likely extremely small. This is also in line with several other studies that were unable to find any effect or meaningful improvements [28].

Importantly, the fact that we did not detect systematic improvements in students’ *grades* does not mean that there were no improvements in *learning*. There is a substantial body of evidence supporting that both acute and chronic sleep loss compromises alertness, cognitive performance and memory, and reduces engagement to perform well (performance effort) [15,62,63]. Thus, improving sleep in sleep-deprived teenagers is very likely to improve their learning [64–66]. In addition, one could speculate that the flexibility and the thus putatively increased self-responsibility and self-determination of students in the flexible system, paired with the reported increase in motivation on later days, may also further improve learning. The question is whether better learning mediated by improved sleep also translates into better grades – and how much sleep improvement is needed and within what timeframe.

Additionally, students’ learning is strongly affected by many factors beyond those captured in our study or those of others on this topic. Models of teaching and learning include several nested factors, such as the individual student (e.g. motivation and prior knowledge), the individual teacher (professional competence [67,68]), the learning environment (e.g. socio-economic status or native language) or factors of instruction (generic and subject-specific instructional quality) but also class-level factors (learning atmosphere or class mates) [69]. Of these, especially instruction and teacher-level factors greatly influence students’ learning [70,71]. Furthermore, grades are inherently suboptimal measures of students’ academic performance, as teachers also include other factors such as compliance, effort, attitude, or behaviour in their assessment [72].

Therefore, it may be a big ask and possibly naive to expect grades to improve noticeably and within a few months after delays or a flexible system have been affected. Rather, we should acknowledge students’ maintained achievements under potentially less effort and improved learning capacities (this needs to be assessed in future studies) in addition to the gift of more sleep and better well-being.

Indeed, teachers at the studied school reported perceiving students as more alert and motivated and tardiness rates as decreased (personal communication). On the other hand, grades still determine future career trajectories and open doors to higher education in many countries [73]. In this sense, they do have a greater importance for careers – at least early on – than other measures of performance, such as standardised or non-standardised tests in class, which might be more valid for measuring academic performance under certain conditions. Additionally, despite all described influences, grades - as indicators of prior knowledge - seem to be the best predictor for achievement in university courses [74].

Our study has several limitations that have not yet been mentioned. We could not obtain information about teachers’ competence, their instructional quality or classroom atmosphere but accounted for gender, quarter, grade level, and discipline - factors that are often overlooked in the field. We also lacked socio-demographic information, which likely influence grades, such as the socioeconomic status (SES), parents’ education or ethnicity of students and their parents [75]. The vast majority of students included in this study were Caucasian by observation, so we had low variation with regards to ethnicity. Lastly, we did not collect objective measures of cognitive performance through cognitive test batteries but asked students to self-evaluate their quality of study [30].

In conclusion, we highlight that current early school start times around the globe are detrimental for sleep and health and likely do not allow students to excel as much as they could. Many studies have shown positive effects on sleep or well-being, when school starts were delayed [21–24] or in systems were school already starts much later, such as in Uruguay or Argentina [76,77]. Thus, it seems fair to argue that later starts are beneficial for students in terms of health and well-being. These factors form a profound basis for good academic achievement but there are also numerous other factors that play into this and possibly mask positive effects: for example, teachers might not perform at their best later during the day or adjust their grading under the new bell times to achieve normal distributions of performance; students might need to spend less time on their homework or learn more easily while still achieving similar but not improved grades; the dose of delay might need to be higher (i.e. more delay or more uptake of later starts in a flexible system) and exposure time might need to be longer until an effect emerges, or grades are insensitive to this kind of intervention, to name only a few. But despite these complications, it should be emphasized that students can maintain their grades *in addition to* better sleep and well-being - a central and very important achievement in its own right. Further studies are required on how to harness the unique advantages of flexible start systems, such as promoting students’ responsibility, choice and investment, for optimal sleep and learning gains.

## Acknowledgements

We thank all the students for their engaged participation and the management of the Gymnasium Alsdorf, in particular W Bock and O Vollert, for their open support throughout the study. We also thank Dr. M Vuori-Brodowski for her contributions during wave 1, general advice and statistical support.

## Author contributions (CRedIT Taxonomy)

Conceptualisation: ECW, TR

Methodology: ECW, AMB

Investigation: AMB, ECW, CM

Data curation: AMB, CM

Formal analysis: AMB, ECW, FO, GZ, CM

Validation: ECW, AMB, FO, GZ

Supervision: ECW

Visualization: AMB, ECW

Writing – original draft: AMB, ECW, CF

Writing – review and editing: AMB, ECW, GZ, FO, CF

## Data availability

Open access sharing of data is not possible due to consent forms that prohibit online deposition of data. We implemented this to secure students’ privacy since students and teachers were well acquainted with each other and might have identified specific participants. Data are available from the corresponding authors upon reasonable request.

## Code availability

We did not develop any custom code or algorithms for data analyses. The code for grade analyses can be found here on github: https://github.com/annambiller/School-study/blob/13bfb38c35332fe285b3cab738b621a4083df7f6/Linear%20mixed%20model%20code%20for%20grade%20analyses.txt

## Supplementary information for

**Figure S1.**
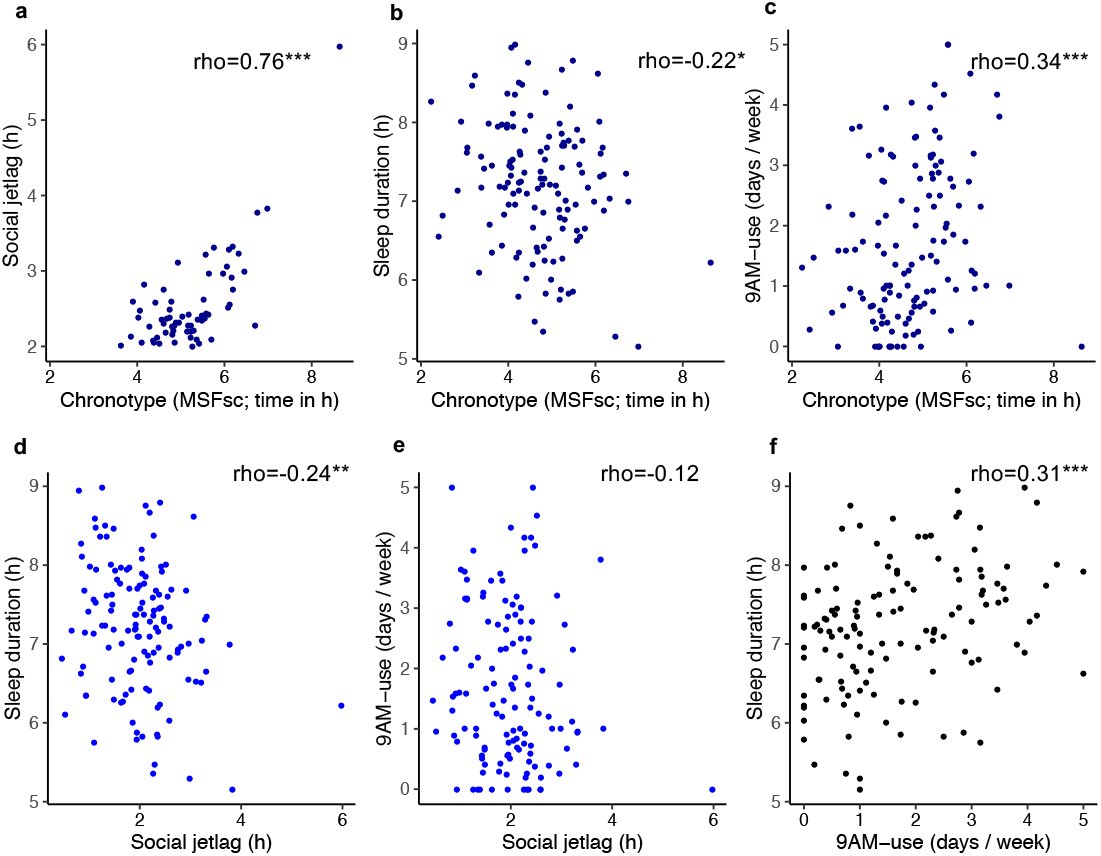
Correlations between sleep variables. Spearman rank correlations between the sleep variables social jetlag, chronotype (MSF_sc_), and sleep duration on schooldays, as well as 9AM-use (frequency of ≥9AM-starts) (n=129). *, p<0.05; **, p<0.01; ***, p<0.001

**Table S1.**
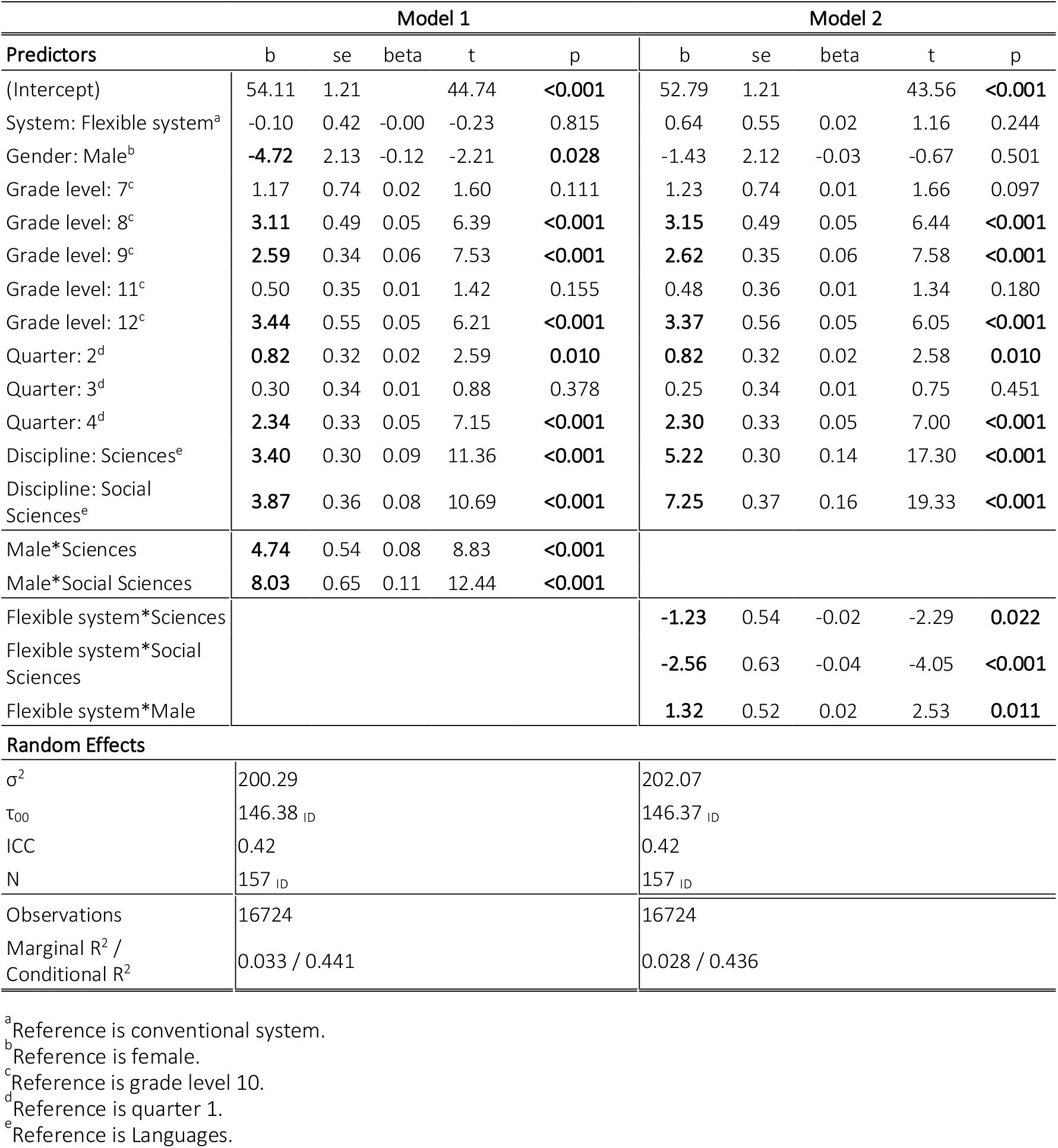
Linear mixed regression models 1 and 2: General and system effects on grades. Predicted outcomes are quarterly grades (0%-100%) in 12 academic subjects from students of cohort 1 and 2 (n=157). Abbreviations: b, unstandardized coefficient; se, standard error; beta, standardized coefficient; t, t-statistic; p, p-value; σ^2^, variance of residuals of random effects; τ_00_, variance of ID intercepts of random effects; ICC, intra-class correlation coefficient (describes how much variance is explained by the random effects); N, number of participants; Marginal R^2^ describes the amount of variance explained by the fixed effects (predictors); Conditional R^2^ describes the amount of variance explained by the full model.

| Model 1 |  |  |  |  |  | Model 2 |  |  |  |  |
| --- | --- | --- | --- | --- | --- | --- | --- | --- | --- | --- |
| Predictors | b | se | beta | t | p | b | se | beta | t | p |
| (Intercept) | 54.11 | 1.21 |  | 44.74 | <0.001 | 52.79 | 1.21 |  | 43.56 | <0.001 |
| System: Flexible system<sup>a</sup> | -0.10 | 0.42 | -0.00 | -0.23 | 0.815 | 0.64 | 0.55 | 0.02 | 1.16 | 0.244 |
| Gender: Male<sup>b</sup> | **-4.72** | 2.13 | -0.12 | -2.21 | **0.028** | -1.43 | 2.12 | -0.03 | -0.67 | 0.501 |
| Grade level: 7<sup>c</sup> | 1.17 | 0.74 | 0.02 | 1.60 | 0.111 | 1.23 | 0.74 | 0.01 | 1.66 | 0.097 |
| Grade level: 8<sup>c</sup> | **3.11** | 0.49 | 0.05 | 6.39 | <0.001 | **3.15** | 0.49 | 0.05 | 6.44 | <0.001 |
| Grade level: 9<sup>c</sup> | **2.59** | 0.34 | 0.06 | 7.53 | <0.001 | **2.62** | 0.35 | 0.06 | 7.58 | <0.001 |
| Grade level: 11<sup>c</sup> | 0.50 | 0.35 | 0.01 | 1.42 | 0.155 | 0.48 | 0.36 | 0.01 | 1.34 | 0.180 |
| Grade level: 12<sup>c</sup> | **3.44** | 0.55 | 0.05 | 6.21 | <0.001 | **3.37** | 0.56 | 0.05 | 6.05 | <0.001 |
| Quarter: 2<sup>d</sup> | **0.82** | 0.32 | 0.02 | 2.59 | **0.010** | **0.82** | 0.32 | 0.02 | 2.58 | **0.010** |
| Quarter: 3<sup>d</sup> | 0.30 | 0.34 | 0.01 | 0.88 | 0.378 | 0.25 | 0.34 | 0.01 | 0.75 | 0.451 |
| Quarter: 4<sup>d</sup> | **2.34** | 0.33 | 0.05 | 7.15 | <0.001 | **2.30** | 0.33 | 0.05 | 7.00 | <0.001 |
| Discipline: Sciences<sup>e</sup> | **3.40** | 0.30 | 0.09 | 11.36 | <0.001 | **5.22** | 0.30 | 0.14 | 17.30 | <0.001 |
| Discipline: Social Sciences<sup>e</sup> | **3.87** | 0.36 | 0.08 | 10.69 | <0.001 | **7.25** | 0.37 | 0.16 | 19.33 | <0.001 |
| Male*Sciences | **4.74** | 0.54 | 0.08 | 8.83 | <0.001 |  |  |  |  |  |
| Male*Social Sciences | **8.03** | 0.65 | 0.11 | 12.44 | <0.001 |  |  |  |  |  |
| Flexible system*Sciences |  |  |  |  |  | **-1.23** | 0.54 | -0.02 | -2.29 | **0.022** |
| Flexible system*Social Sciences |  |  |  |  |  | **-2.56** | 0.63 | -0.04 | -4.05 | <0.001 |
| Flexible system*Male |  |  |  |  |  | **1.32** | 0.52 | 0.02 | 2.53 | **0.011** |
| **Random Effects** |  |  |  |  |  |  |  |  |  |  |
| $\sigma^2$ | 200.29 | | | | | 202.07 | | | | |
| $\tau_{00}$ | 146.38<sub>ID</sub> | | | | | 146.37<sub>ID</sub> | | | | |
| ICC | 0.42 |  |  |  |  | 0.42 |  |  |  |  |
| N | 157<sub>ID</sub> |  |  |  |  | 157<sub>ID</sub> |  |  |  |  |
| Observations | 16724 |  |  |  |  | 16724 |  |  |  |  |
| Marginal R<sup>2</sup> / Conditional R<sup>2</sup> | 0.033 / 0.441 |  |  |  |  | 0.028 / 0.436 |  |  |  |  |

<sup>a</sup>Reference is conventional system.
<sup>b</sup>Reference is female.
<sup>c</sup>Reference is grade level 10.
<sup>d</sup>Reference is quarter 1.
<sup>e</sup>Reference is Languages.

**Table S2.** Post hoc results of mixed models 1 and 2. Results are presented as marginal estimated means of quarterly grades scaled 0-100% (standard error), degrees of freedom. Simple contrast results are presented as estimated difference of academic grades (standard error), p-value. Degrees of freedom method: Kenward-Rogers. Results are averaged over the levels of system or gender, grade level, and quarter. Tukey method for comparison of 3 estimates.

| Model 1 |  |  |  |  |
| --- | --- | --- | --- | --- |
| Gender | Languages | Sciences | Social Sciences | Simple contrasts |
| female | 56.7 (1.19)<br>166 | 60.1 (1.19)<br>166 | 60.6 (1.21)<br>176 | Languages-Sciences:<br>**-3.41 (0.30), p<.0001**<br>Languages-Social Sciences:<br>**-3.87 (0.36), p<.0001**<br>Sciences-Social Sciences:<br>**-0.47 (0.36), p=0.3895** |
| male | 52.0 (1.79)<br>164 | 60.2 (1.78)<br>163 | 63.9 (1.80)<br>172 | Languages-Sciences:<br>**-8.15 (0.45), p<.0001**<br>Languages-Social Sciences:<br>**-11.90 (0.54), p<.0001**<br>Sciences-Social Sciences:<br>**-3.76 (0.52), p<.0001** |
| Simple contrasts | 4.72 (2.13)<br>p=0.0284 | -0.00 (2.13)<br>p=0.9915 | -3.31 (2.16)<br>p=0.1269 |  |
| Model 2 |  |  |  |  |
| System | Languages | Sciences | Social Sciences | Simple contrasts |
| conventional | 54.7 (1.08)<br>168 | 59.9 (1.08)<br>167 | 62.0 (1.10)<br>182 | Languages-Sciences:<br>**-5.22 (0.30), p<0.0001**<br>Languages-Social Sciences:<br>**-7.25 (0.38), p<0.0001**<br>Sciences-Social Sciences:<br>**-2.03 (0.37), p<0.0001** |
| flexible | 56.0 (1.14)<br>212 | 60.0 (1.13)<br>206 | 60.7 (1.16)<br>227 | Languages-Sciences:<br>**-3.99 (0.45), p<.0001**<br>Languages-Social Sciences:<br>**-4.69 (0.51), p<.0001**<br>Sciences-Social Sciences:<br>**-0.69 (0.49), p=0.3355** |
| Simple contrasts | -1.30 (0.54)<br>p=0.0168 | -0.07 (0.61)<br>p=0.8849 | 1.26 (0.61)<br>p=0.0384 |  |
| System | Female | Male |  | Simple contrasts |
| conventional | 59.6 (1.18)<br>160 | 58.2 (1.77)<br>158 |  | female-male:<br>1.43 (2.12), p=0.5010 |
| flexible | 59.0 (1.22) | 58.9 (1.81) |  | female-male: |
|  | 186 | 173 |  | 0.11 (2.14), p=0.9580 |
| **Simple contrasts** | 0.62 (0.45)<br>p=0.1726 | 0.7 (0.56)<br>p=0.2106 |  |  |

**Table S3.**
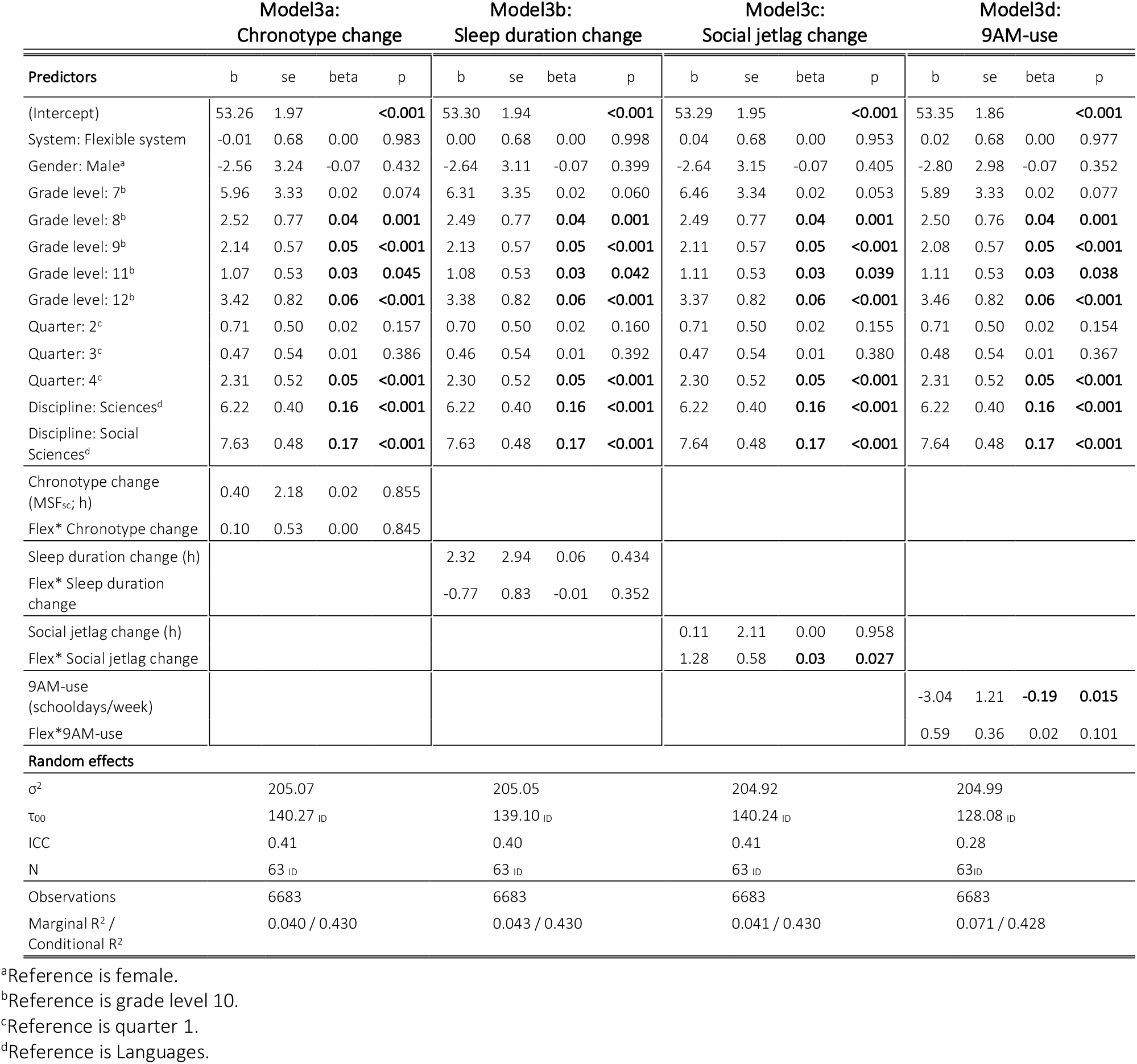
Linear mixed regression models 3a-d: Effect of changes in sleep and of ≥9AM-use on grade improvements from the conventional to the flexible system. Predicted outcomes are quarterly grades (0%-100%) in 12 academic subjects from students of cohort 1 (n=63) over 4 years. “Change” refers to the absolute difference of the respective sleep variable between the conventional and the flexible system (t1-t0). Positive numbers mean later chronotype, longer sleep and more social jetlag in the flexible system (t1). 9AM-use is the frequency of ≥9AM-starts at t1 (no baseline data for calculation of change available). Abbreviations: Flex, Flexible system; b, unstandardized coefficient; se, standard error; beta, standardized coefficient; p, p value; σ^2^, variance of residuals of random effects; τ_00_, variance of ID intercepts of random effects; ICC, intra-class correlation coefficient (describes how much variance is explained by the random effects); N, number of participants; Marginal R^2^ describes the amount of variance explained by the fixed effects (predictors); Conditional R^2^ describes the amount of variance explained by the full model.

**Table S4.** Linear mixed regression models 4a-e: Effect of absolute sleep characteristics in the flexible system on grades. Predicted outcomes are quarterly grades (0%-100%) in 12 academic subjects from students of cohorts 1 and 2 (n=129) over 1.5 years in the flexible system. 9AM-use is the frequency of ≥9AM-starts in the flexible system. Abbreviations: b, unstandardized coefficient; se, standard error; beta, standardized coefficient; p, p value; σ^2^, variance of residuals of random effects; τ_00_, variance of ID intercepts of random effects; ICC, intra-class correlation coefficient (describes how much variance is explained by the random effects); N, number of participants; Marginal R^2^ describes the amount of variance explained by the fixed effects (predictors); Conditional R^2^ describes the amount of variance explained by the full model.

|  | Model 4a:<br>Chronotype |  |  |  | Model 4b:<br>Sleep duration |  |  |  | Model 4c:<br>Social jetlag |  |  |  | Model 4d:<br>9AM-use |  |  |  | Model 4e:<br>All |  |  |  |
| --- | --- | --- | --- | --- | --- | --- | --- | --- | --- | --- | --- | --- | --- | --- | --- | --- | --- | --- | --- | --- |
| Predictors | b | se | beta | p | b | se | beta | p | b | se | beta | p | b | se | beta | p | b | se | beta | p |
| (Intercept) | 59.50 | 5.81 |  | **<0.001** | 65.13 | 10.70 |  | **<0.001** | 53.68 | 3.40 |  | **<0.001** | 60.50 | 2.19 |  | **<0.001** | 64.77 | 14.51 |  | **<0.001** |
| Gender: Male<sup>a</sup> | -0.31 | 2.61 | -0.01 | 0.907 | -0.61 | 2.54 | -0.01 | 0.810 | -0.83 | 2.54 | -0.02 | 0.744 | -0.72 | 2.49 | -0.02 | 0.773 | -0.15 | 2.63 | 0.00 | 0.956 |
| Grade level: 9<sup>b</sup> | 3.35 | 0.82 | **0.05** | **<0.001** | 3.35 | 0.82 | **0.05** | **<0.001** | 3.36 | 0.82 | **0.05** | **<0.001** | 3.27 | 0.82 | **0.05** | **<0.001** | 3.27 | 0.82 | **0.05** | **<0.001** |
| Grade level: 11<sup>b</sup> | 0.47 | 0.62 | 0.01 | 0.450 | 0.46 | 0.62 | 0.01 | 0.451 | 0.45 | 0.62 | 0.01 | 0.469 | 0.58 | 0.62 | 0.01 | 0.350 | 0.57 | 0.62 | 0.01 | 0.355 |
| Grade level: 12<sup>b</sup> | 0.02 | 1.00 | 0.00 | 0.984 | 0.02 | 0.99 | 0.00 | 0.984 | -0.02 | 0.99 | 0.00 | 0.983 | 0.27 | 1.00 | 0.01 | 0.790 | 0.26 | 1.00 | 0.01 | 0.794 |
| Quarter: 2<sup>c</sup> | 0.38 | 0.65 | 0.01 | 0.559 | 0.38 | 0.65 | 0.01 | 0.559 | 0.38 | 0.65 | 0.01 | 0.559 | 0.38 | 0.65 | 0.01 | 0.562 | 0.38 | 0.65 | 0.01 | 0.561 |
| Quarter: 3<sup>c</sup> | -0.05 | 0.65 | -0.00 | 0.933 | -0.06 | 0.65 | -0.00 | 0.932 | -0.07 | 0.65 | 0.00 | 0.917 | 0.02 | 0.65 | 0.00 | 0.980 | 0.02 | 0.65 | 0.00 | 0.979 |
| Quarter: 4<sup>c</sup> | 1.69 | 0.61 | **0.04** | **0.005** | 1.69 | 0.61 | **0.04** | **0.005** | 1.68 | 0.61 | **0.04** | **0.006** | 1.76 | 0.61 | **0.05** | **0.004** | 1.76 | 0.61 | **0.05** | **0.004** |
| Discipline: Sciences<sup>d</sup> | 4.27 | 0.43 | **0.11** | **<0.001** | 4.28 | 0.43 | **0.11** | **<0.001** | 4.27 | 0.43 | **0.11** | **<0.001** | 4.27 | 0.43 | **0.11** | **<0.001** | 4.28 | 0.43 | **0.11** | **<0.001** |
| Discipline: Social Sciences<sup>d</sup> | 4.65 | 0.49 | **0.10** | **<0.001** | 4.65 | 0.49 | **0.10** | **<0.001** | 4.65 | 0.49 | **0.10** | **<0.001** | 4.65 | 0.49 | **0.10** | **<0.001** | 4.65 | 0.49 | **0.10** | **<0.001** |
| Chronotype (local time in h) | -0.53 | 1.23 | -0.03 | 0.665 |  |  |  |  |  |  |  |  |  |  |  |  | -2.37 | 2.35 | -0.12 | 0.315 |
| Sleep duration (h) |  |  |  |  | -1.12 | 1.47 | -0.05 | 0.448 |  |  |  |  |  |  |  |  | -0.27 | 1.61 | -0.01 | 0.865 |
| Social jetlag (h) |  |  |  |  |  |  |  |  | 1.76 | 1.54 | 0.07 | 0.256 |  |  |  |  | 3.70 | 2.78 | 0.14 | 0.186 |
| ≥9AM-use (schooldays/week) |  |  |  |  |  |  |  |  |  |  |  |  | -2.12 | 0.91 |  | **0.022** | -1.32 | 1.20 | -0.08 | 0.272 |
| **Random Effects** |  |  |  |  |  |  |  |  |  |  |  |  |  |  |  |  |  |  |  |  |
| $\sigma^2$ | 168.88 | | | | 168.87 | | | | 168.87 | | | | 168.87 | | | | 168.87 | | | |
| $\tau_{00}$ | 174.27<sub>ID</sub> | | | | 173.89<sub>ID</sub> | | | | 172.93<sub>ID</sub> | | | | 167.17<sub>ID</sub> | | | | 168.81<sub>ID</sub> | | | |
| ICC | 0.51 |  |  |  | 0.51 |  |  |  | 0.51 |  |  |  | 0.50 |  |  |  | 0.50 |  |  |  |
| N | 129<sub>ID</sub> |  |  |  | 129<sub>ID</sub> |  |  |  | 129<sub>ID</sub> |  |  |  | 129<sub>ID</sub> |  |  |  | 129<sub>ID</sub> |  |  |  |
| Observations | 5111 |  |  |  | 5111 |  |  |  | 5111 |  |  |  | 5111 |  |  |  | 5111 |  |  |  |
| Marginal R<sup>2</sup> / Conditional R<sup>2</sup> | 0.018 / 0.517 |  |  |  | 0.019 / 0.517 |  |  |  | 0.022 / 0.517 |  |  |  | 0.037 / 0.516 |  |  |  | 0.042 / 0.521 |  |  |  |

<sup>a</sup>Reference is female.
<sup>b</sup>Reference is grade level 10.
<sup>c</sup>Reference is quarter 1.
<sup>d</sup>Reference is Languages.

